# Who is researching biodiversity hotspots in Eastern Europe? A case study on grasslands from Romania

**DOI:** 10.1101/487397

**Authors:** Andreea Nita, Tibor Hartel, Steluta Manolache, Cristiana M. Ciocanea, Iulia V. Miu, Laurentiu Rozylowicz

## Abstract

Farming landscapes of Europe are vital arenas for social-ecological sustainability because of their significant coverage and potential to integrate food production with biodiversity conservation. Knowledge gathered by scientific research is a critical ingredient for developing and implementing socio-economically and ecologically sustainable grassland management strategies for grasslands. The quality of scientific knowledge and its potential to address grasslands as complex social-ecological systems is strongly dependent on the creativity and scientific ambition of the researcher, but also on the network (from academic and non-academic sectors) around the researcher. The goal of this paper is to map the research network around Romania’s grasslands. These systems have exceptional socio-cultural and economic values and are between the most biodiverse ecosystems of the world. Considering the multiple threats to these grasslands, it is an urgent need to understand the existing scientific knowledge profile around these systems. This paper aims at using bibliometrics analysis, a well-developed scientific domain that envisages network theory to analyze relationships between affiliations network, co-authorship network, and co-word analysis. The number of studies targeting grassland management in Romania is increasing mainly thanks to international involvement. However, the management of the grasslands is still deficient and the contribution of science to the process is virtually absent. The subject of research is mainly related to the biological and ecological characteristics of grasslands, a notable absence from internationally visible research being the management of grasslands, especially in the context of EU Common Agricultural Policies. To increase scientific performance, and better inform EU and local policies on grassland management, Romanian researchers should better capitalize on international collaborations and local academic leaders. Our findings can be used to identify research gaps and to improve collaboration and knowledge exchange between practitioners, scientists, policy makers, and stakeholders.

## Introduction

The integration of agricultural production, biodiversity conservation, and socio-cultural values is a key challenge for the sustainability of social-ecological systems (Fischer et al. 2012). Current grasslands of Europe developed under millennia-long human management, typically by livestock grazing and hay production. Grasslands have a substantial contribution to the high nature value farmlands o the European Union (Veen et al. 2009, Plieninger and Bieling 2013, Lomba et al. 2014). Several protected species and habitats are linked to grasslands and are dependent on some form of extensive, multifunctional management of these ecosystems (Halada et al. 2011, Dorresteijn et al. 2017). Furthermore, wooded meadows and wood-pastures are considered as an archetypical manifestation of HNV farmlands in Europe (Plieninger et al. 2015).

Many EU countries already lost culturally and naturally important grasslands due to changes in management practices (Pe’er et al. 2014). The intensive, highly specialized management of the grasslands resulted in the sharp decrease in their biodiversity, aesthetic and cultural values or even the disappearance of the grasslands. On the other hand, land abandonment also threatens several species and habitats (however also creating opportunities for conserving others) as well as ecosystem services of grasslands, commonly through woody vegetation encroachment (Stoate et al. 2009, Bugalho et al. 2011, Queiroz et al. 2014, Michielsen et al. 2017).

Knowledge is essential for a socioeconomically and ecologically sustainable grassland management (Fischer et al. 2012). In one hand, scientific knowledge can generate a contextual understanding of the relationship between the management intensity and the biodiversity and productivity of the grasslands (Kleijn et al. 2009). On the other hand, scientific research is important in developing new types of conceptualizations of the grasslands, for example as complex, adaptive systems which can exist in multiple social-ecological states (Sutcliffe et al. 2014, Hartel et al. 2018). Nevertheless, scientific knowledge can contribute to re-addressing current management paradigms around the management of production landscapes (Abson et al. 2017), including grasslands. Since grasslands can simultaneously fulfill multiple social-ecological values and roles (e.g., production, biodiversity conservation, recreational and cultural values), an overly narrow (e.g., disciplinary) scientific approach for understanding them could promote simplistic management measures which often lack socio-cultural contextualization. The quality of scientific knowledge and its potential to address grasslands as complex social-ecological systems is strongly dependent on the creativity and scientific ambition of the researcher, but also on the social network (from academic and non-academic sectors) around the researcher (Lescourret et al. 2015).

In this study, we address the collaboration network between the academics and the diversity of research domains around the Romanian grasslands. The importance of our research is fourfold. First, Romanian grasslands are between the best biodiversity hotspots at the global level (Wilson et al. 2012), with several ancient land-use forms such as wood-pastures (Cremene et al. 2005, Plieninger et al. 2015), traditional stewardship and management forms (Babai and Molnár 2014). These grasslands are under threat from overgrazing, changes in management and stewardship and abandonment (Baur et al. 2006, Ruprecht 2006, Sutcliffe et al. 2014, Peringer et al. 2017). Second, Romania is a developing country in Eastern Europe, where research is suffering from lack of funds, institutional instability and intense political pressure (David and Marko 2018, Miclaus and Micu 2018). This overall harsh conjuncture for research is hampering the production of holistic knowledge (Hanspach et al. 2014) and innovation which will be indispensable for Romania in order to navigate challenges of globalization, including its increasing role in the global food security (Benton et al. 2011), coping with extreme climate variations (Azadi et al. 2018) and land grabbing (Petrescu-Mag et al. 2017). Third, the academic world in Romania is still transitioning from a local/regional, disciplinary thinking and approaches towards adopting international standards of scientific rigor and holistic, inter- and transdisciplinary approaches. Because of this, Romania is sharply underrepresented in the international scientific databases while publishing in local journals (i.e., hosted by the academic institutions) is still actively promoted by academic institutions. Fourth, management of grasslands in Romania is still deficient (Pătru-Stupariu et al. 2017) despite the latest legal motions, which most often do not consider the contribution of science to the process.

Collaboration between academics and knowledge creation have been extensively studied using network analysis framework. Analyses span from understanding of patterns of scientific collaboration as resulted from co-authorship and institutional networks (Newman 2004, Hancean et al. 2014, Hancean and Perc 2016) to knowledge creation (Wang 2016), to prediction of productivity (Barabási et al. 2002) or trends (Chen et al. 2010). In this paper we use network analysis, to characterize the status of scientific research in the field of grassland governance in Romania and suggests ways to improve scientific performance by: 1) revealing internationally visible research around Romania’s grasslands published after 1990; 2) highlighting most important institutions generating the research, and mapping the invisible authors and academic leaders as resulted from the co-authorship network, and 3) analyzing the co-occurrence keyword network to discover the most common keywords, research topics, and scientific interest.

## Methods

To identify the research related to grasslands in Romania, we extracted from the Scopus database (Elsevier B.V.) 602 articles, book chapters, and conference proceedings potentially related to the investigated subject. We obtained these publications by searching simultaneously abstracts, titles, and keywords sections with the following keywords: *Common Agricultural Policy, CAP, pasture, grassland, meadow, lawn, greensward, grazing, graze, silvopastoral, pastureland, rangeland, mowing* (and adding *Romania* to each of them, e.g., “pasture” AND “Romania”). Then, we went through each of these publications and removed those who do not contain information about the subject of our review (e.g., paleoecology, paleobotany) and published before 1990. In this way, we obtained a final database that includes 197 publications relevant to the grasslands of Romania (Figure 1). For each article, we extracted the list of keywords, authors and their affiliations as stated in the papers. Figure 1 shows an ascending trend in the number of publications, which can be interpreted as an indicator of increasing interest of scientific researchers on grasslands from Romania

**Figure 1.**
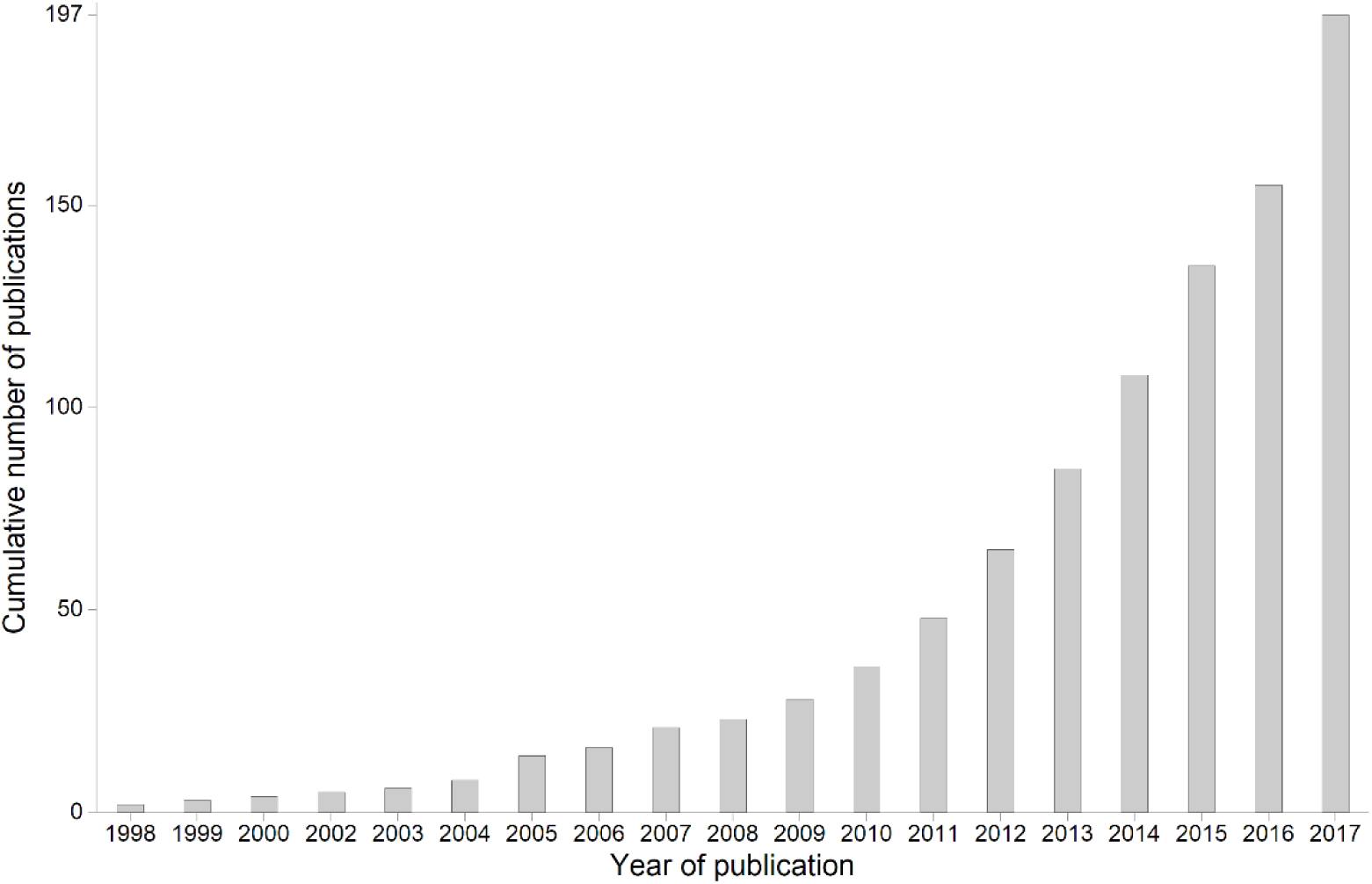
Cumulative number of publications targeting Romania’s grasslands accessible on the Scopus database (1998-2017).

The scientific articles targeting Romanian grasslands were published in 107 journals and proceedings. Top journals in our network, with more than 5 publications are: *Quality-Access to Success* (15 publications), *International Multidisciplinary Scientific GeoConference Surveying Geology and Mining Ecology Management* (SGEM) (14), *Notulae Botanicae Horti Agrobotanici Cluj-Napoca* (8), *PLoS ONE* (6), *Applied Vegetation Science* (5), *Biodiversity and Conservation* (5), *Environmental Engineering and Management Journal* (5), and *North-Western Journal of Zoology* (5).

For the bibliometric analysis, we constructed three network matrices: (i) an affiliations network, to infer about inter-institutional cooperation; (ii) a co-authorship network, which highlight invisible authors and academic leaders, and (iii) and a co-occurrence keyword network to discover hidden connections between the most common keywords, research topics and scientific interest. For these analyses, we created distinct databases which we then cleaned and unified (i.e., standardize the name of institutions, authors, and keywords to avoid duplication of entries due to different spelling). The cleaned matrices include 192 unique affiliations (516 entries initially), 517 unique authors (755 entries initially) and 577 unique keywords (1019 entries initially). The methodology workflow is presented in Figure 2.

**Figure 2.**
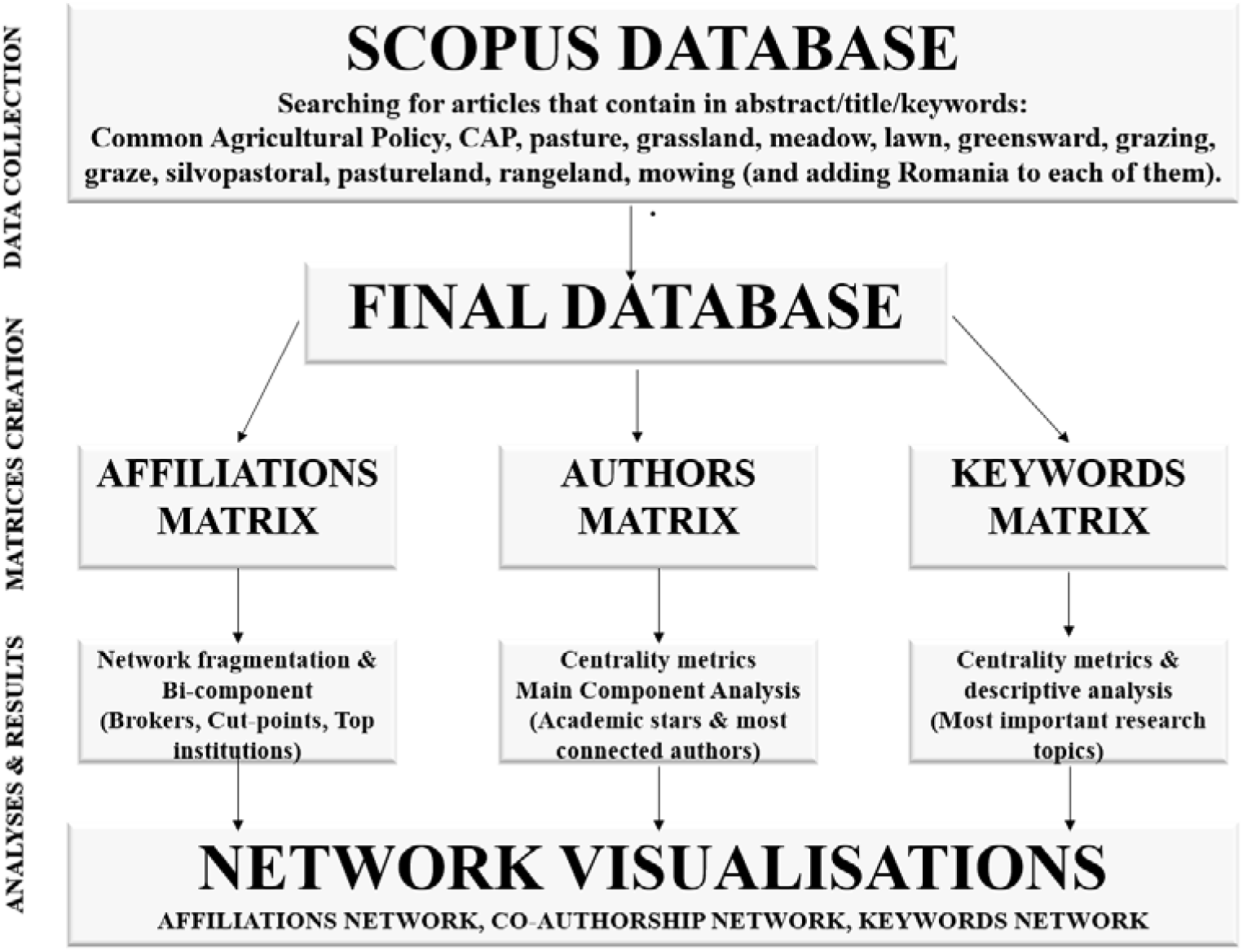
Workflow of network analyses of internationally visible research related to grasslands of Romania.

### (i) Affiliations analysis

We used *bi-component analysis* to identity blocks (bi-connected subnetworks) and the cut-points (articulation points) in the affiliation graph (Hanneman and Riddle 2005). If removed, cut point institutions break the affiliation network into one or more bi-connected subnetworks (Leydesdorff 2004). Such institutions are important for cohesivity of research network focused on Romanian pastures and might act as research brokers among otherwise disconnected groups (Borgatti et al. 2018). An affiliation subgraph is bi-connected if every institution in the subgraph (minimum three) has direct connections to other and even removing any node, the subgraph remains connected (Hanneman and Riddle 2005). The number and size of blocks is an indication of network fragmentation. If a network is dominated by one block, then most institutions cooperate during the research. Smaller blocks include isolated institutions, which published a small number of papers on the subject or on uncommon topics (Leydesdorff 2004).

We also calculated several node-level centrality metrics: degree, betweenness, and eigenvector. *Degree centrality* of an institution represents the number of direct connections (Abbasi et al. 2012, Borgatti et al. 2018). The metric identifies the most collaborative institutions, i.e., the institutions with the highest number of connections to other research units in Romanian grassland research. The metric does not account for how important the institutions are linked to the node of interest. Thus, an institution can be considered as collaborative even if it is linked only to institutions that have no other collaborations in the network. *Eigenvector centrality* is similar to degree centrality but scores higher connections with institutions which are themselves well connected (Borgatti et al. 2018). It represents the sum of the eigenvectors of the institutions that the institution of interest is connected to, and reveal the best options for future partnerships (most influent institutions, which can further promote partnership in research) (Borgatti et al. 2018). *Betweennesses centrality* measures the extent to which an institution lies on paths between other institutions from the research network. Such institutions can control the flow of information in the network if we assume that every pair of connected institutions exchanges information with equal probability and information flow on a short path chosen at random (Barabási 2016). Such institutions can be seen as bridge affiliations, e.g., can control the research subjects in a partnership (Leydesdorff 2004). Relationships among countries and within countries, resulted from affiliations declared by the authors, were represented using a chord diagram (Flor 2018).

### (ii) Authors analysis

We use authors’ matrix to analyze the co-authorship network created around Romanian grassland research and to illustrate the patterns of cooperation between the scientists in this research domain (Barabási et al. 2002, Ding et al. 2014). We calculated the *network fragmentation* metric, which in our case indicates the proportion of pairs of authors that cannot reach each other (Borgatti et al. 2018). Large datasets, as it is the case of our database, typically involve many small independent clusters around larger ones and one large and dense cluster (Hancean and Perc 2016). Thus, we assessed the distribution of components in the co-authorship network and extracted the largest connected component (main component) that shows the group of highly active authors focused on grasslands at the national level (subnetworks of authors that are maximally connected between each other). We also, calculated the *degree centrality* in order to find out which are the most collaborative authors within the network, and compared the results with the *betweenness centrality* to highlight the “academic stars” within the network – authors with high degree and betweenness centralities (Glänzel and Schubert 2004, Ding et al. 2014).

To better understand the motivations for scientific collaboration the Romanian grassland research, three authors with key positions in the research network were asked to respond to an interview on the opportunities and constraints of research in this domain. A network leader affiliated to a Romanian institution (*network leader a*), a foreign author continuing scientific work in Romania (*network leader b*), and a foreign author who works abroad (*network leader c*),

### (iii) Keywords network

Analyses based on keywords have been applied in various techniques such as text mining, data reduction and clustering (Cobo et al. 2011, Ding et al. 2014, Popescu et al. 2014) to identify emerging research. Keywords are also good representatives for the main topics addressed by research in general (Dotsika and Watkins 2017), and co-occurrence *keywords network* analysis can be used to highlight the most common and important research keywords (Ding et al. 2014). Hence, to infer about most common, popular, and bridge keywords featured in Romanian grasslands research, we used the degree, eigenvector, and betweenness centralities (see affiliations networks for details about these network-level metrics).

Network analyses were performed in UCINET software (Borgatti et al. 2002), and the networks were graphically represented using Netdraw (Borgatti 2002), and *chorddiag* R package (Flor 2018).

## Results

### Affiliations network

The network of organizations hosting researchers publishing about grasslands from Romania includes 192 distinct institutions from 36 countries out of which 14 are isolated institutions (i.e. only collaborates inside their institution). Nine of the isolated affiliations are from foreign countries (Japan, Poland, Italy, Spain, Germany, Switzerland, Ukraine), but there are also 5 isolated institutions from Romania: Politehnica University of Timisoara, University of Pitesti, Technical University of Cluj-Napoca, University Bogdan Voda, and Danube Delta National Institute for Research and Development. The best-represented affiliations involved in Romanian grassland research are from Romania (53), Germany (23), United Kingdom (13) and Hungary (13).

Researchers from Romanian institutions collaborate mostly with researchers from Romania (138 institutional collaborations), Germany (92), Hungary (29), Czech Republic (19), and UK (18). Also, the next best-represented country, Germany, collaborate mostly with institutions from Romania and Germany (Figure 3).

**Figure 3.**
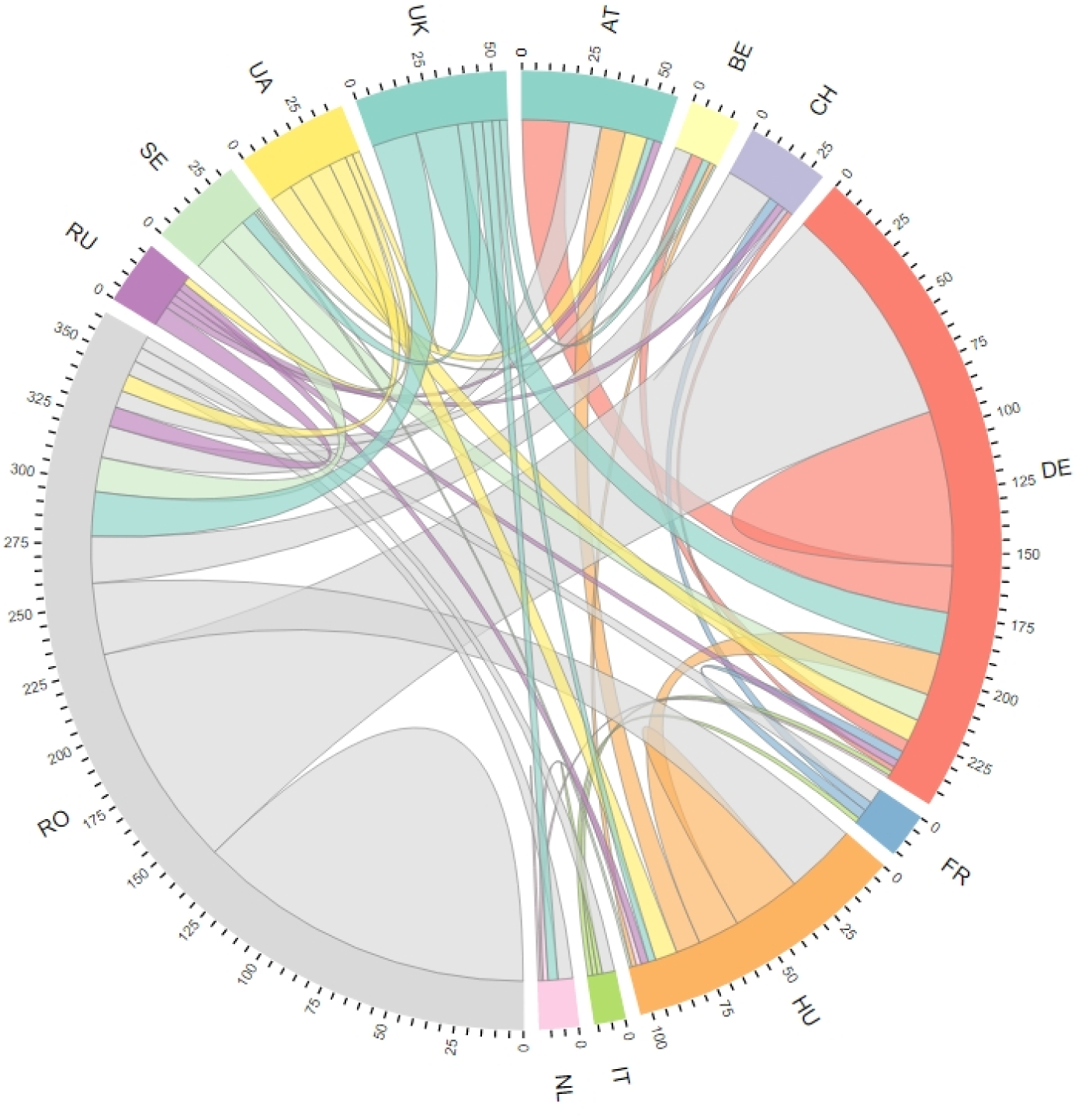
Chord diagram showing international and national collaborative pattern as resulted from internationally visible publications related to grasslands in Romania (links = co-occurring countries in an article; only countries with > 10 collaborations with institutions from Romania are shown).

The Bi-components analysis generated 45 blocks (Supplementary Table 1) held by 25 cut points/affiliation brokers (Figure 4), which if they were to be removed, the structure of the network would become divided into unconnected parts. Block 31 has the highest number of affiliations (101, out of which 20 are from Romania). Institutions acting as affiliation brokers are from Romania (16 institutions), Hungary (3), Slovenia (1), Czech Republic (1), Germany (2), Sweden (1) and Italy (1).

**Figure 4.**
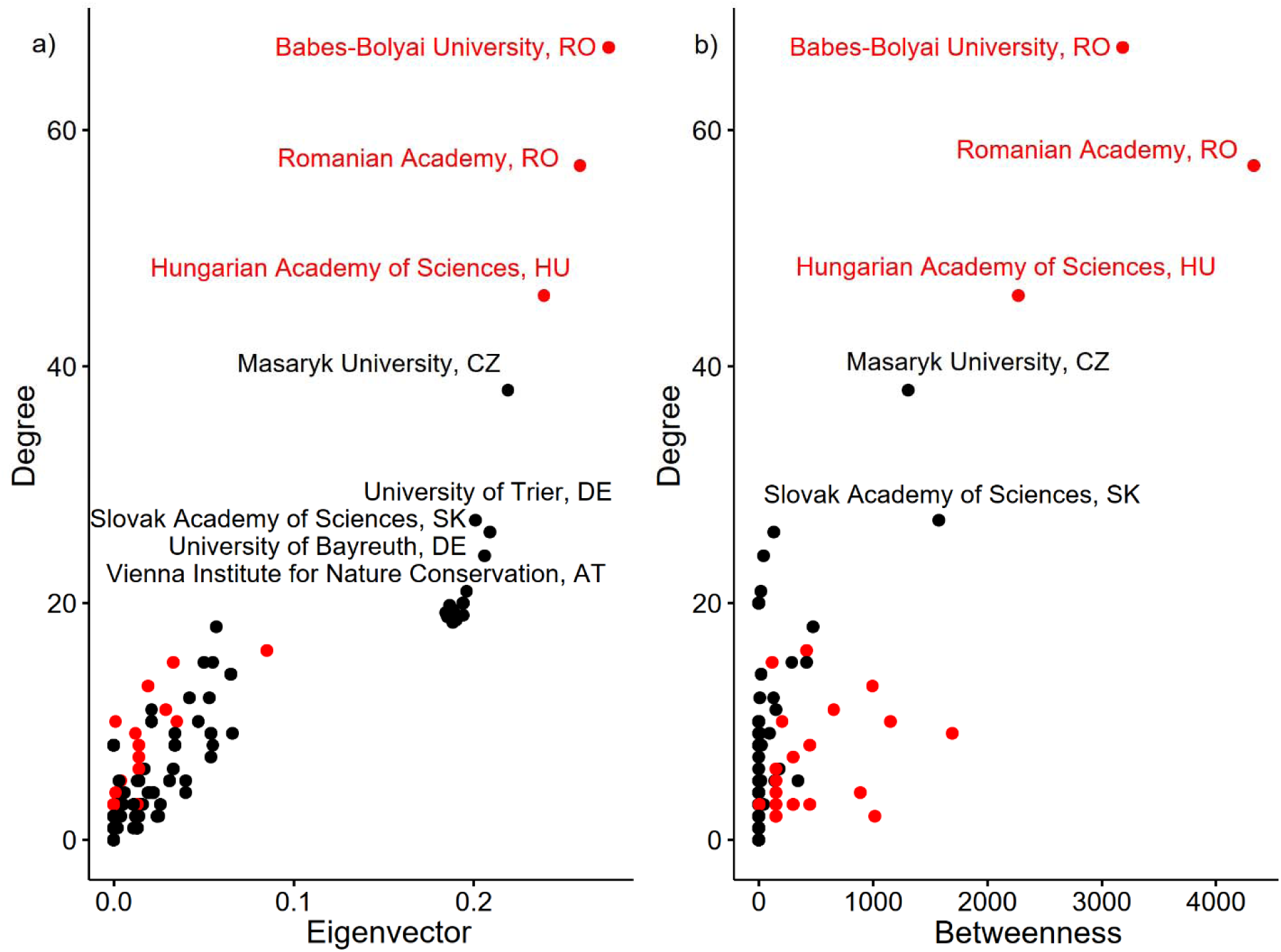
Degree versus eigenvector (a) and betweenness (b) centralities of institution hosting researchers publishing about Romania’s grasslands (cut points/affiliation brokers are in red).

Figure 4 presents the affiliations ranked by their importance in terms of best position within the bibliometric network (betweennesses centrality), influence (eigenvector centrality), and number of network connections relevant for Romanian grassland research (degree centrality). The *Babes-Bolyai University* (Romania) control the flow of information (betweenness), has the highest number of connections (degree), and is the most influent institution (eigenvector) within the network (Supplementary Table 1). The *Romanian Academy* (Romania) also has the control over the entire network from the scientific cooperation point of view and occupies the second position in term of number of connections and influence (Figure 4 and 5) while on the 3^rd^ position is the *Hungarian Academy of Sciences* (Hungary).

**Figure 5.**
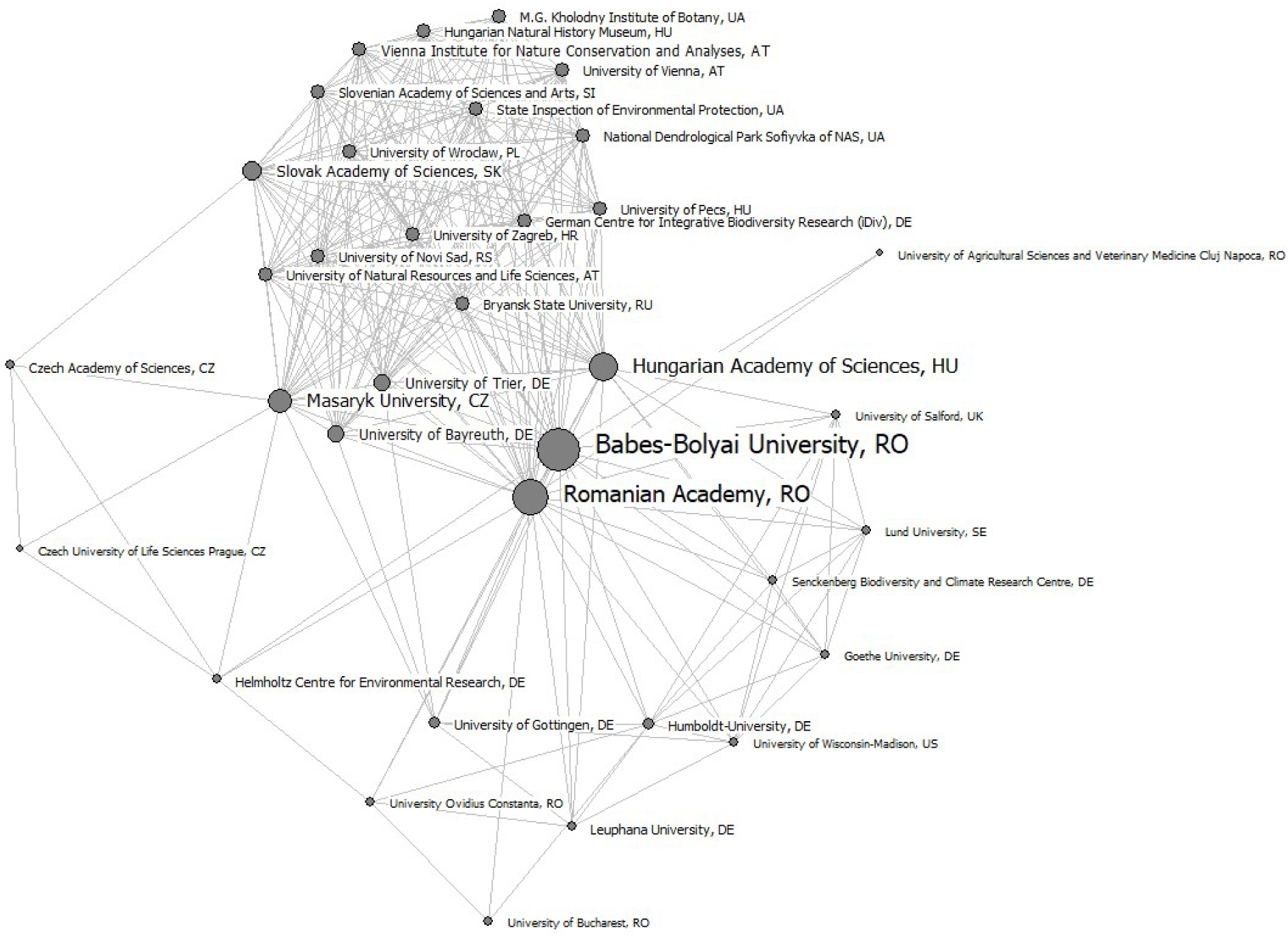
Network of the affiliations of authors publishing about Romania’s grasslands with a degree centrality >10 (size of nodes and labels given by degree).

### Authors network

The network of authors is composed of 517 authors and has a high fragmentation (0.886). The main component analysis divided the fragmented network into 92 clusters (Supplementary Table 1). Cluster 6 is the densest component and sums up 168 authors (Figure 6), while the rest of the clusters have a mean of 3.83 authors (stdev = 3.97). Giving its importance, we further mapped Cluster 6 to have a closer look at its structure (Figure 6). Most of the authors forming part of the main component are not Romanians, hence, many important authors within the Romanian grassland research network are foreigners (Figure 7).

**Figure 6.**
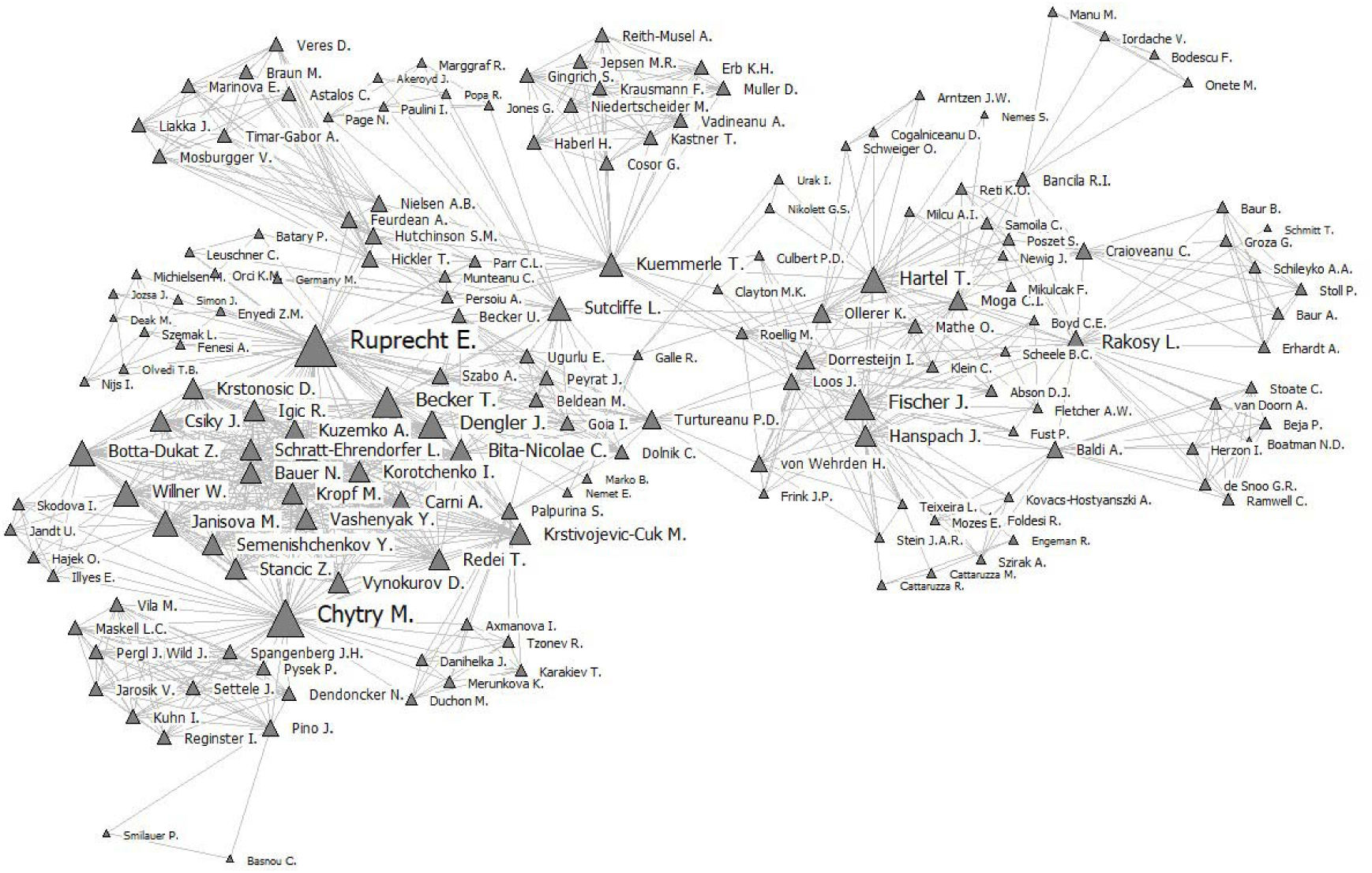
Main component of the network of authors publishing about Romania’s grasslands (size of nodes given by degree).

**Figure 7.**
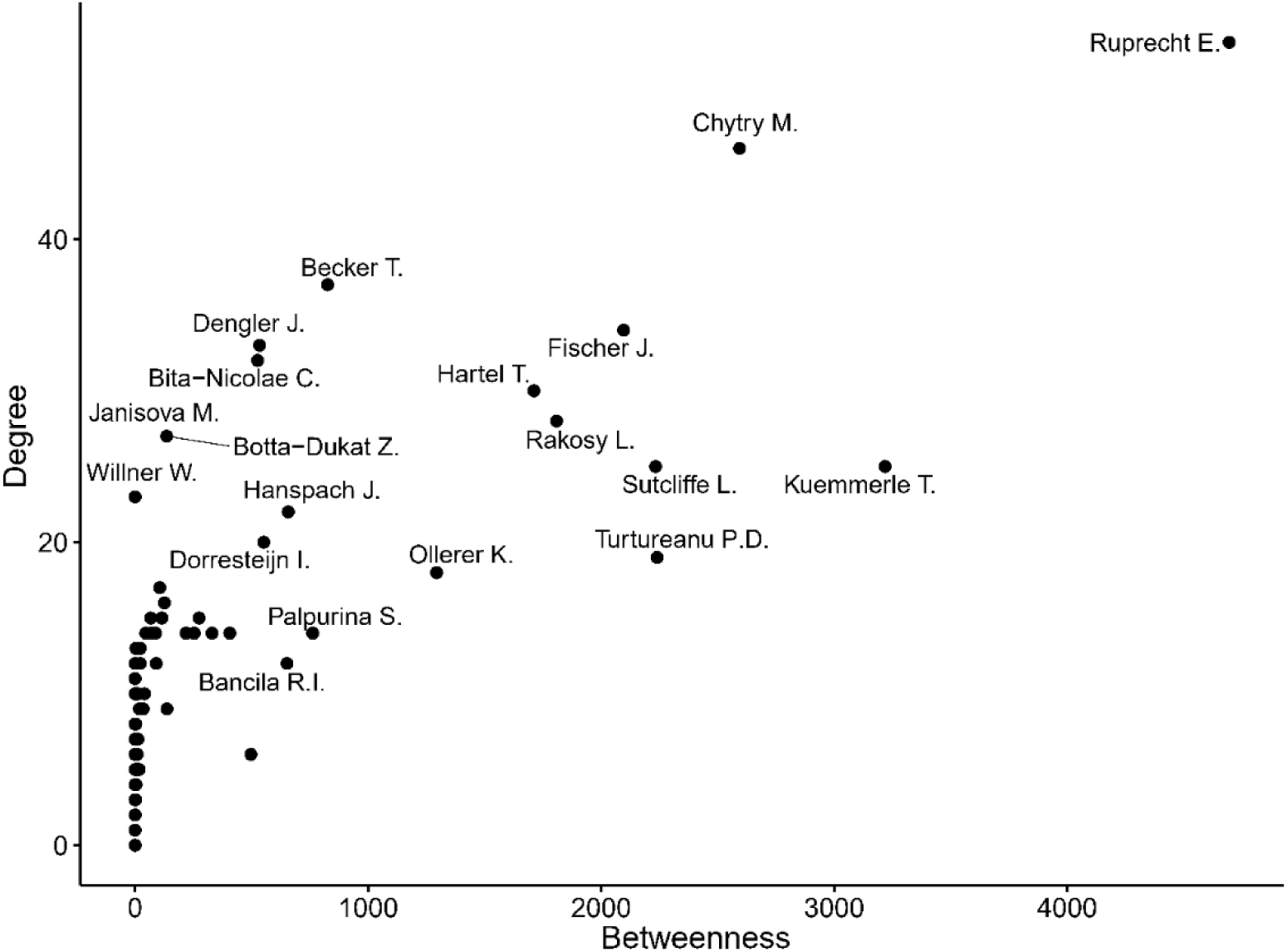
Top academic authors publishing about Romania’s grasslands in terms of number of connections (degree) and position in the network (betweenness).

Based on this result, we carried out a short interview with three network leaders in order to conclude how they have evolved in their professional career, who inspired them to research the grasslands of Romania and what is the key to their success in terms of national and international partnerships made in this field. The authors were anonymized as “*network leader a”* (NLa), “*network leader b*” (NLb), and “*network leader c*” (NLc).

The paragraph below summarizes the answers of interviewed network leaders for the following questions.

**Q1. Who inspired you to research grasslands in Romania (e.g., researchers, articles, institutions from Romania or foreign).**

**NLa:** *After finishing my studies at the Babe*□*-Bolyai University from Cluj-Napoca, Romania, I have met a Romanian grassland researcher with a great expertise and a passion for rare plants of steppe habitats. Field trips to beautiful steppe-like grasslands and the concern about their transformation or loss inspired me to study grasslands. Later on, my Ph.D. supervisor, especially his way of thinking about diversity, inspired me a lot, thus I have worked with him and some of his colleagues for several years.*

**NLb:** *A friend was researching High Nature Value Farmland in Europe and told me about the species-rich grasslands in Transylvania that she had visited. I was curious, and with her assistance and the help of local partners, I was lucky enough to be able to do some fieldwork for a scientific study there.*

**NLc:** *It was in 1997 before I studied Biology when I first came to Transylvania. I stayed for a few days with some friends in a small village not far from Sighisoara. It was summer and one evening we climbed a hill behind the houses to watch the sunset. I don’t remember the sunset, but I do remember the meadows we walked through. They were so amazing. Although I have grown up on a farm myself, I hadn’t seen nor smelled nor heard such a meadow before. Every plant seemed to be blooming and every insect buzzing. It was such richness and beauty. That was the thing that made me most enthusiastic about grasslands in Transylvania, even if I only later learned to put scientific labels on that richness.*

**Q2. How did you form your co-authorship or partnership network? What were the main challenges and opportunities in this respect?**

**NLa:** *At the very beginning of my career, personal contacts have been the most important in finding mentors, since there was a limited access to scientific literature and even to the internet in my home country in the 1990s. These first mentors gave me opportunities to further develop my ‘partnership network’ by inviting me to their institutions abroad (in Hungary, where I had no language or cultural barriers) and to conferences, where I could meet other scientists. Later on, I have contacted scientists from abroad (outside of Hungary), whose papers raised my interest the most, by e-mail, and looked for scholarships for research stays at their host institutions. Nowadays, there are scientists from European institutions contacting me and asking me for collaboration within the framework of scientific projects.*

**NLb:** *The Romanian conservation NGO Foundation ADEPT generously hosted my work in Romania, and through them, I met many, many interesting people involved in all aspects of conservation (academic and non-academic). Perhaps the most difficult thing was to stay in touch with all of them!*

**NLc:** *My network developed through a social-ecological research project which included research on vascular plant diversity in the farming landscape of Southern Transylvania (Sighisoara). That means through multiple pathways including cooperation with local NGOs, scientists, field assistants, and friends. Opportunities clearly included to learn from knowledge and experiences from others and to join skills for getting fieldwork and analysis done. As a challenge, I see the language barrier and also that we didn’t do a classic vegetation study but applied a random sampling design for putting survey sites (which didn’t match with the more traditional vegetation research).*

**Q3. How would you describe the collaboration between Romanian grassland researchers?**

**NLa:** *There are few collaborations, and the majority is based on personal acquaintance. Confidence or convenience (people working at the same institution) have important roles in forming research teams around a certain project.*

**NLb:** *I only have experience of my own collaboration with Romanian grassland researchers, which was pleasant and productive.*

**NLc:** *To be honest, I don’t feel I can say much about this. I don’t have much insight into their collaboration. Also, I don’t consider myself a grassland expert and therefore I am not so deep into this network.*

**Q4. How would you describe the role of foreign scientists in Romanian grassland research?**

**NLa:** *Foreign scientists find Romanian grasslands remarkable, and they come to work in Romania with very specific research questions. They often look for a local expert in Romanian grasslands to involve in their project, and by this means collaborations arise. I consider foreign scientists bring in many interesting research questions, achieve nice results and invigorate Romanian grassland research.*

**NLb:** *My impression is that scientists from northern and western Europe are very aware of the grassland diversity that has been lost in the more intensified regions of Europe. I think in any ecosystem there is always value in combining observations from local researchers who intimately know the area and can interpret its subtleties, and “outsiders” who can maybe bring a fresh perspective and spot parallels with other systems.*

**NLc:** *From my limited perspective they seem to have a strong influence or at least they seem to stand out for me. I could name a few foreign grassland researchers but would struggle to name the same number of Romanians. However, this is probably biased because I am a foreigner myself, I might pay more attention to fellow foreigners that are active in this field.*

### Keywords network

To find the most researched topics focused on grasslands, we mapped the keyword network after excluding the occasionally used keywords, that is, those who had a degree of less than 10 (Figure 8). Not considering the words that were used to search for the articles that were used for creating the study database, the keywords most influent and important within the network were *Biodiversity* and *Conservation* (Figures 8, 9). Also, our results showed that terms such as *Farm management, Pastoral value, Landscape pattern, Ancient trees* are among many other keywords on the bottom positions, both from the perspective of their use and their importance within the network (Supplementary Table 1).

**Figure 8.**
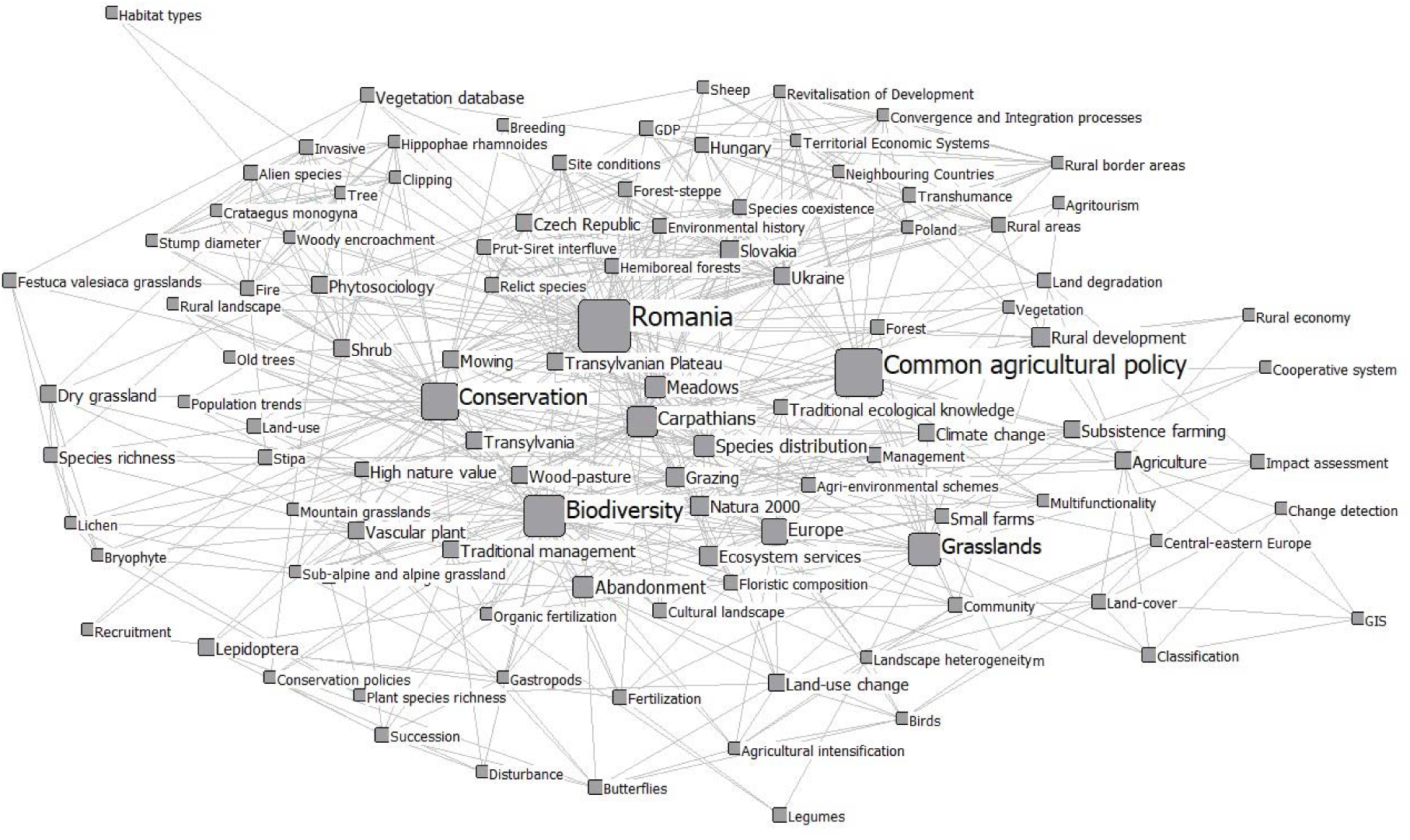
Top keywords used in publications about Romania’s grasslands by degree (size of labels and nodes by degree, for a better visualization we illustrate only the keywords with a degree > 10).

**Figure 9.**
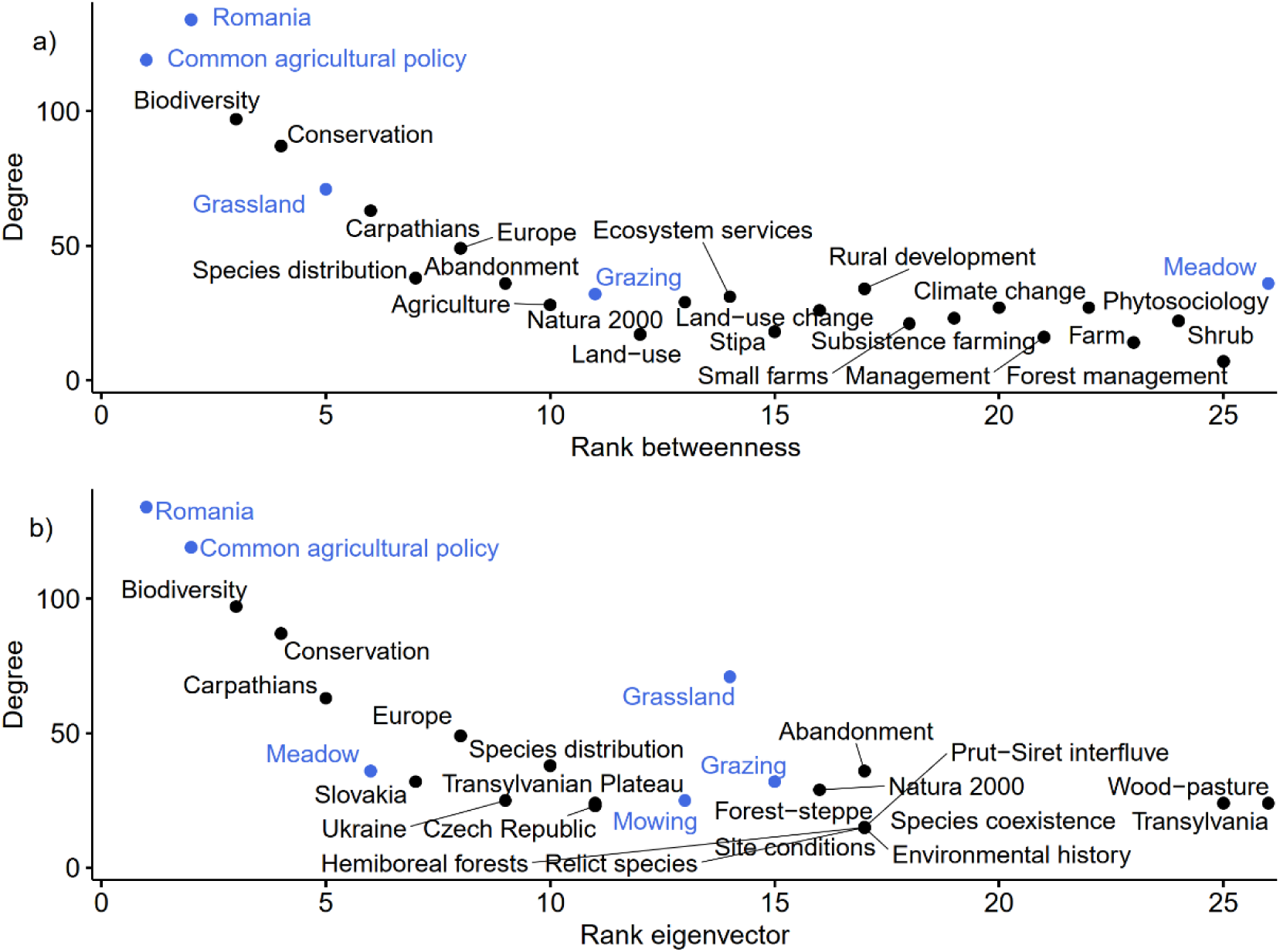
Top 25 keywords used in publications about Romania’s grasslands by degree versus a) betweenness and b) eigenvector centralities (blue = keywords used for data collection).

## Discussion

We provide a comprehensive analysis of the research network around grasslands in an Eastern European country, Romania. Grasslands have key socio-cultural, economic and natural values and can be approached from several disciplinary perspectives (Veen et al. 2009). Thus, research on grasslands are important contributions to the knowledge because of the several perspectives within which these systems can be understood, including agronomy, nature conservation, ecology, sociology, statistics, sociology, anthropology, economics, political sciences, geography. We found that both national and international researchers and institutions contribute to scientific knowledge regarding Romanian grasslands, with a higher influence from foreign researchers. Furthermore, we identified the most commonly addressed and the most influential research topics regarding Romanian grasslands.

### Who is researching Romanian grasslands?

Considering the limited number of articles, our result shows a great international coverage of institutions and researchers involved in Romanian grassland research (192 institutions, 517 coauthors, 197 articles). By analyzing the co-authorship centrality metrics, we showed that the contribution of foreign institutions and researchers to grasslands research is high, while there are also key institutions from Romanian which have an important role as brokers. We are aware of the fact that a single author can name several host institutions in the research papers, however, first nominated institution usually is the home institution (Glänzel and Schubert 2004) which reduce the potential bias. This redundancy (i.e., one author feeling a responsibility to indicate multiple institutional addresses) also may represent opportunities for innovative knowledge generation which can flow into the cumulative knowledge pool (Leydesdorff 2004, Newman 2004). For example, one of the authors of the present manuscript (TH) adopted a social-ecological approach for understanding wood-pasture systems of Romania and Europe; this approach was adopted as a result of a postdoctoral research period spent at Leuphana University (Lüneburg, Germany), where the research team addressed a holistic understanding of the sustainability challenges of cultural landscapes in Romania.

We found that a researcher can have an outstanding contribution to the knowledge of Romanian grasslands not only as ‘grassland specialist’ but also as an ecological modeler, trans-disciplinary researcher or economist (see also the interviews for the profile of three researchers highlighted by our analysis), as being part of a larger, interdisciplinary research group. While internationally visible scientific production sharply increased to a maximum of 42 papers in 2017, relatively few papers are published in top-tier journals, as in other Eastern European countries (Kozak et al. 2015). The lack of improvement in the publishing performance can be explained by the limited funding (David and Marko 2018, Miclaus and Micu 2018) as well as by the established traditions in choosing the target journals (Campos-Arceiz et al. 2015). Inter-disciplinary and international expertise is an important driver of the knowledge generation (Gazni et al. 2012) for grasslands management, and researchers of Romania could capitalize on this even more. International collaborations are increasingly possible and encouraged, e.g., within Horizon 2020 and Biodiversa-EraNet project partnerships (Granieri and Renda 2012). These collaborations can allow knowledge flow, the development of new coauthorship networks (Hancean and Perc 2016) and also can buffer the unstable funding which characterizes Romanian research (Miclaus and Micu 2018). Our results suggest several ways in which a local researcher can increase its attractivity for international research projects, including a keen interest in a holistic understanding of the cultural landscapes (of which grasslands are part), active and long-term engagement with local stakeholders and partners and increased scientific productivity (Balvanera et al. 2017). The interviews of the academic stars of the network show, irrespective of the origin of the author that answered, that the cooperation to investigate Romanian grasslands is made more by means of recommendations and by common knowledge, the role of foreign authors and institutions being of defining importance regarding research initiatives and partnerships.

### What were the research topics addressing Romanian grasslands?

We identified 577 keywords in internationally visible research addressing Romanian grasslands. Few keywords have high importance in the overall keyword network; this can be interpreted as the main topics driving grassland research in Romania. These keywords (Figures 8, 9) are mostly related to the high natural values of grasslands, ecosystem services and the land-use practices (including abandonment) related to grasslands. The results can be explained in multiple ways.

First, there was a momentum generated by the accession of Romania to the European Union (2007), where the delineation of Natura 2000 sites and the development of management plans for them was and are still an ongoing process (Manolache et al. 2017). Second, the establishment of conservation biology as a research discipline in the academic environment of Romania (besides the classical ecology research, especially in the 2000s, also resulted in research projects which targeted rare species and habitats as well as the negative impact of management (especially overgrazing) on these. Third, several Non-Governmental Organizations also increased the social awareness about the decline of biodiversity (especially in the protected areas but also beyond) (Rozylowicz et al. 2017), this again motivating research targeting the biodiversity and conservation of grasslands. Within this, the mountain hay meadows have outstanding importance (this is why ‘Carpathians’ were highlighted as important, Figures 8 and 9), being highly biodiverse as well as threatened by overgrazing and abandonment (Cremene et al. 2005, Sutcliffe et al. 2014, Michielsen et al. 2017). A relatively recent overview of the research targeting Natura 2000 sites in the European Union showed that research supporting Natura 2000 network is dominated by ecological research while the policy and social aspects are underrepresented (Popescu et al. 2014). Our analysis on Romanian grasslands research suggests a similar pattern. We identified several research topics which we would expect to be better represented (i.e., more influential) in the network, because of their crucial importance in the management of the grasslands. Less important topics from the perspective of network analysis are the economy of grasslands, the traditional ecological knowledge related to grasslands, stakeholders and the non-herbaceous elements across the grasslands. From the perspective of holistic understanding (Hanspach et al. 2014), there are recently established research projects in different regions of Romania (i.e., ‘Sustainable Landscapes in Central Romania’), which aim to contribute to a socially and ecologically sustainable farming system in Romania. Romania hosts some of the most representative wood-pasture systems of Central and Eastern Europe (Roellig et al. 2018). Despite their common occurrence in Romania and their relative scarcity in Western Europe (Plieninger et al. 2015), we found a surprisingly low number of papers addressing these systems. Since wood-pastures of Romania suffered from the lack of tree regeneration (Roellig et al. 2018) and the erosion of values related to scattered trees (Torralba et al. 2018), it is of utmost importance to amplify the holistic research on these systems. Based on our collective, long-term experience as researchers within the Romanian academic system, we believe that Romanian academia still has much to do for implementing holistic, trans-disciplinary research to address sustainability problems related to farming landscapes in general and pastures in particular. One major barrier of adopting an integrative approach is the strong tradition of disciplinary research which still dominates research and teaching at the universities. This is also reflected in the dominant research themes identified in Figures 8 and 9.

## Conclusions

Although grasslands are complex social-ecological systems which can be studied in several scientific domains or interdisciplinary, internationally visible research networks around Romania’s grasslands is still undeveloped (e.g., relatively low number of papers in top-tier, low number of visible researchers with institutional affiliation from Romania). The co-authorship network structure reveals several institutional leaders who can further promote the research in this area. These top institutions are prestigious institutions from Romania closely followed by foreign collaborators (e.g., from Hungary, Germany). Based on their academic profile, top researchers are from diverse scientific fields (plant ecology, conservation biology, population ecology, etc.), a feature favoring the scientific performance by increasing the interdisciplinary and relevancy of research. The subject of research is mainly related to the biological and ecological characteristics of grasslands, a notable absence from internationally visible research being the management of grasslands, especially in the context of EU Common Agricultural Policies. To increase scientific performance, and better inform EU and local policies on grassland management, Romanian researchers should better capitalize on international collaborations and local academic leaders.

## Supporting information

## Acknowledgments

This work was supported by a grant of the Romanian National Authority for Scientific Research (https://uefiscdi.ro/), PN-III-P4-IDPCE-2016-0483. We would like to thank the three network leaders for their valuable contribution added to our paper through their interview answers.

